# Initial and ongoing tobacco smoking elicits vascular damage and distinct inflammatory response linked to neurodegeneration

**DOI:** 10.1101/2022.09.27.509833

**Authors:** Alejandra P. Garza, Lorena Morton, Éva Pállinger, Edit I. Buzás, Stefanie Schreiber, Björn H. Schott, Ildiko Rita Dunay

**Affiliations:** Institute of Inflammation and Neurodegeneration, Medical Faculty, Otto-von-Guericke University Magdeburg, Germany; Department of Genetics, Cell- and Immunobiology, Semmelweis University, Budapest; HCEMM SU Extracellular Vesicle Research Group, Budapest; ELKH-SE Translational Extracellular Vesicle Research Group, Budapest; Department of Neurology, Otto-von-Guericke University, Magdeburg, Germany; Center for Behavioral Brain Sciences, Magdeburg, Germany; German Center for Neurodegenerative Diseases (DZNE), Magdeburg, Germany; Leibniz Institute of Neurobiology, Magdeburg, Germany; Department of Psychiatry and Psychotherapy, University Medicine Göttingen, Göttingen, Germany

**Keywords:** Tobacco smoking, innate immunity, extracellular vesicles, monocytes, vascular damage, neurodegeneration

## Abstract

Tobacco smoking is strongly linked to vascular damage contributing to the development of hypertension, atherosclerosis, as well as an increased risk for neurodegeneration. Still, the contribution of the innate immune system to the development of vascular damage upon chronic tobacco use before the onset of clinical symptoms is not fully elucidated. Notably, our data provide evidence that a single acute exposure to tobacco in never smokers elicits a secretion of extracellular vesicles by endothelial cells expressing CD105 and CD49e, granting further recognition of early preclinical biomarker of vascular damage. Further, we investigated the effects of smoking on the immune system of healthy asymptomatic chronic smokers compared to never-smokers and focused on the innate immune system. Our data reveal a distinct immune landscape representative for early stages of vascular damage before tobacco smoking related disease develop in clinically asymptomatic chronic smokers. These results indicate a dysregulated immuno-vascular axis in chronic tobacco smokers that are considered healthy individuals. The distinct alterations are characterized by increased CD36 expression by blood monocyte subsets, neutrophilia, increased plasma IL-18 and reduced levels of IL-33, IL-10 and IL-8. Further, the detection of lower circulating BDNF and elevated sTREM2, specific markers for neurodegeneration, suggests a considerable pre-clinical impact of tobacco smoking on CNS function in clinically healthy individuals. These findings provide further insight into the initial and ongoing effects of tobacco smoking and the potential vascular damage contributing to the progression of neurodegenerative disorders, specifically cerebrovascular dysfunction and dementia.

## Introduction

Tobacco smoking is a major modifiable risk factor for a plethora of pathological conditions, and the number of smokers continues to rise globally, despite largely undisputed adverse health effects (Gowing et al. 2015; Eriksen M et al. 2012). Currently, 22.3% of the world population smoke, with detrimental effects on both life expectancy and quality of life (Cheng and Jin 2022). It has been predicted that half of the smoker population will die prematurely from tobacco-related complications (World Health Organization 2021). Diverse systemic chronic disorders are directly attributable to continuous exposure to cigarette smoke, with the lungs and cardiovascular system most severely affected, but also involving the central and peripheral nervous systems (Yanbaeva et al. 2007). In aging societies, the role of smoking as a modifiable risk factor for dementia is of particular importance. Tobacco smoking is a major risk factor for vascular dementia (O’Brien and Thomas 2015), most likely through toxic effects of cigarette contents resulting in vascular damage (Livingston et al. 2020), and meta-analytic evidence further points to an increased risk for Alzheimer’s disease (AD) in smokers (Anstey et al. 2007).

A likely pathomechanism mediating the negative impact of cigarette smoking on vascular and brain health may be through cigarette smoke promoting systemic inflammatory processes and thereby endothelial damage. The constant and chronic presence of tobacco smoke as a stressor, unequivocally leads to inflammation and endothelial dysfunction (Golbidi et al. 2020). Tobacco smoking is strongly linked to vascular endothelial damage, thus contributing to the development of hypertension and atherosclerosis (Messner and Bernhard 2014). Cigarette smoke induces a series of mechanisms that activate cell populations from both the innate and the adaptive immunity, which in turn promote the secretion of multiple inflammation-related molecules such as proinflammatory cytokines including chemokines, reactive oxygen species (ROS) and extracellular vesicles (EVs) (Miteva et al. 2018; Elisia et al. 2020; Kodidela et al. 2020).

Alterations in both the innate and the adaptive immunity have previously been described in the context of tobacco smoking (Miteva et al. 2018; Piaggeschi et al. 2021; Delgado et al. 2021). T cell activation, differentiation and response are altered in chronic smokers, and previous evidence *in vitro* and *in vivo* points to an increase in cell percentages, particularly in CD4+T helper subsets, accompanied by an increased release of proinflammatory cytokines (Forsslund et al. 2014; Zhang et al. 2014). Regarding innate immunity, previous studies have described elevated levels of circulating blood leukocytes in chronic smokers, including neutrophils and monocytes, as well as an increase of peripheral inflammatory markers such as fibrinogen, C-reactive protein (CRP) and proinflammatory cytokines like IL-6 and IL-1β (Andersson et al. 2019; Elisia et al. 2020; Pedersen et al. 2019). *In vitro* studies have demonstrated that after the addition of cigarette smoke extract (CSE), THP-1 cells, a widely used monocytic cell line, become activated along with the nucleotide oligomerization domain (NOD), leucine-rich repeat (LRR), containing pyrin domain-3 (NLRP3) inflammasome, the activation of which leads to the production of IL-1β and IL-18 (Mehta and Dhawan 2020; Qin 2012; Swanson et al. 2019). However, the explicit role of immune cell activation in the pathophysiology of endothelial damage and atherosclerosis has not been fully elucidated. One focus of our study is to understand the role of the peripheral monocytes expressing CD36, especially at the initiation of endothelial damage. CD36 is a scavenger receptor expressed on the surface of monocytes, which mediates uptake of oxidized LDL (oxLDL) from the lumen and thus, facilitates the conversion of healthy peripheral monocytes into foam cells that initiate the development of vascular plaques (Febbraio et al. 2001; Park 2014). CD36 has also been described as a potential early biomarker of atherosclerosis due to its role in plaque formation, angiogenesis and endothelial inflammation (Tian et al. 2020).

Tobacco smoke activates a number of intracellular signaling pathways (Lee et al. 2008; Hellermann et al. 2002; Baskara et al. 2020). However, smoke-induced EV release after a single session of tobacco smoking in never-smokers has not yet been described. EVs are nanosized membrane-enclosed particles derived from all types of cells in the organism. They are released via membrane budding or exocytosis under steady state conditions, during cell activation or apoptosis and play an important role in both cellular homeostasis and cell to cell signaling (Buzas et al. 2014; Osteikoetxea et al. 2016; Tóth et al. 2021). Specifically, EVs have not been investigated extensively *in vivo* in the context of smoking populations. *In vitro* studies have shown however, that EVs are released from alveolar macrophages in response to tobacco smoke and that cultivated bronchial epithelial cells increase the production of EV-miRNAs that contribute to lung carcinogenesis (Li et al. 2010; Liu et al. 2016).

A considerable body of evidence demonstrates that oxidative stress has a detrimental role and promotes dysfunction of different barriers including the blood-brain barrier (BBB) (Hossain et al. 2009; Sajja et al. 2015; Hawkins et al. 2004). In chronic smokers, the increased ROS production may promote endothelial dysfunction predisposing to morphological and functional damage of the BBB, thereby contributing to the pathogenesis of cerebrovascular and neurodegenerative diseases (Zalba et al. 2007; Sophocles Chrissobolis et al. 2011; Pietro Scicchitano et al. 2019; Cataldo et al. 2010). Changes in cardiovascular fitness, with consequent changes in the endothelium and BBB can potentially lead to neurovascular alterations (Cipollini et al. 2019; O’Brien and Thomas 2015). By understanding the initial steps of this process, we aimed to detect early alterations on the neuroinflammation response, to prevent downstream pathologies.

Soluble triggering receptor expressed on myeloid cells 2 (sTREM2) is the secreted domain of TREM2, a receptor expressed on microglial cells, and it has been described as a microglial activation marker and constitutes a promising neurodegeneration biomarker in both CSF and peripheral blood of AD patients (Ferri et al. 2020; Hu et al. 2014; Bekris et al. 2018; Park et al. 2021). Notably, recent evidence points to a gene-environment interaction of smoking and a TREM2 genetic variation contributing to AD risk (Sun et al. 2022). Conversely, with respect to neuronal integrity, plasticity, and survival, brain-derived neurotrophic factor (BDNF) constitutes an established biomarker. BDNF can cross the BBB, and therefore, BDNF levels measured in the periphery correlate with those found in the brain (Pan et al. 1998; Klein et al. 2011; Zuccato and Cattaneo 2009). Results regarding peripheral BDNF levels in smokers vary with some studies reporting reduced plasma concentrations (Bhang et al. 2010), while others detected increased plasma BDNF levels in smokers (Galle et al. 2021).

Here we demonstrate the early and ongoing effects of tobacco smoking and related vascular damage in healthy individuals, that might contribute to the development of cerebrovascular dysfunction and related dementia. Our study indicates the rapid release of EVs into blood circulation after an acute tobacco smoking intervention by the damaged endothelium. Neutrophilia and an elevated number of blood monocytes were detected in chronic smokers. Monocyte subsets increased the expression of CD36, a scavenger receptor involved in atherosclerotic plaque formation. Importantly, decreased levels of brain-derived neurotrophic factor (BDNF) and an elevated sTREM2 in plasma levels in smokers indicate that smoking contributes to the progression of neurodegenerative disorders in otherwise heathy individuals.

## Methods

### Participants

The present study prospectively enrolled 76 participants designated as current smokers (n=26) and nonsmokers (n=50) (Table 1). Participants were recruited from June to August 2020 via advertisement at the Otto-von-Guericke University Campus in Magdeburg, Germany. All enrolled patients were ≥18 years of age, able to give written informed consent and had no cardiovascular, neurological or psychiatric diseases. They reported negatively to current or recent history of infections, tumors, immunizations, autoimmune disease and pregnancy. Participants had no history of uncontrolled metabolic, respiratory or systemic disorders, and current or recent history of either centrally acting medication (e.g., anticonvulsants, antidepressants, anxiolytics, neuroleptics, opioids), anti-inflammatory pharmacotherapy (e.g., non-steroidal anti-inflammatory drugs (NSAIDs), antibiotics, corticosteroids), or anticoagulants. Smoking status was self-reported and defined as never or current smokers. Never smokers are subjects who never smoked or have smoked less than 100 cigarettes in their lifetime but never on a daily basis. Current smokers are otherwise healthy participants who have smoked more than 100 tobacco cigarettes in their lifetime and currently smoke daily. Subjects were asked to arrive in fasting conditions and to refrain from smoking for at least 8 hours. Out of the nonsmokers, 20 participants took part in the acute intervention study (Figure 1A). The acute setting consisted of blood withdrawal before and 60 minutes after smoking two cigarettes (12 tar yield mg/cig, 1.0 nicotine yield). Our study was conducted in accordance with the ethical principles for the Declaration of Helsinki with approval from the ethics committee of the Medical Faculty, Otto-von-Guericke Magdeburg (No 07/17; addendum 12/2021). Written informed consent was obtained from all participants enrolled in this study.

**Figure 1.**
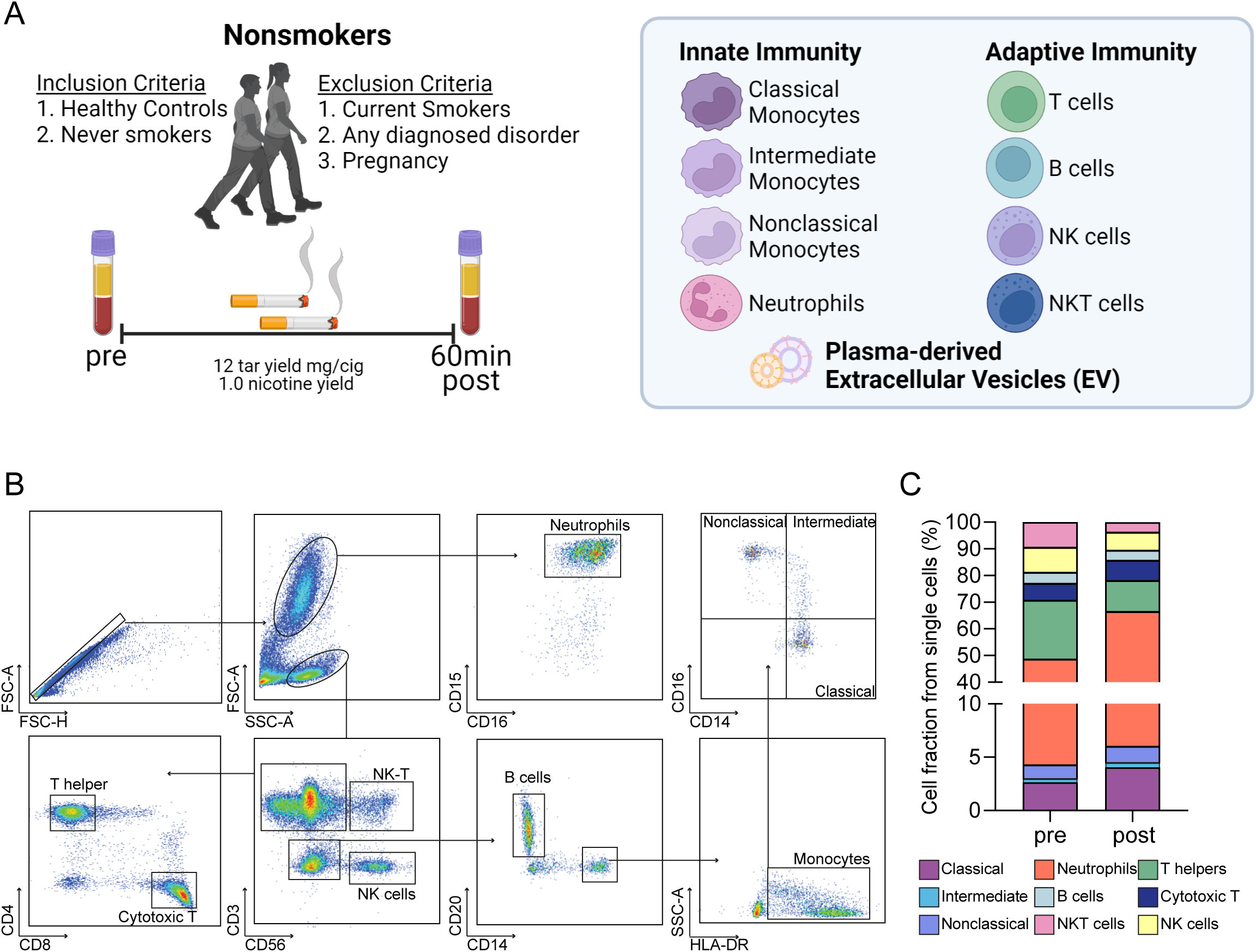
Subtle shift in the innate and adaptive immune cells in healthy controls after one tobacco smoking session. Clinical and experimental study design and analytical processes showing inclusion and exclusion criteria for nonsmoking healthy controls. Blood was withdrawn before and 60 minutes after the second and last cigarette consumption and further analysis of immune system cell composition and extracellular vesicles was performed (**A**). Representative gating strategy to identify granulocytes and mononuclear cells based on their size and granularity. Neutrophils were gated based on surface expression of CD16 and CD15. Mononuclear cells were gated and further identified as T cells, NK cells and NKT cells. The expression of CD4 and CD8 was further assessed in the CD3^+^ T cells. B cells were defined as CD3^-^, CD56^-^ and CD20^+^ cells. Monocytes were identified as CD3^-^, CD56^-^ and CD20^+^, and HLA-DR_+_ and CD14^+^. Gated monocytes were further divided into subpopulations based on their differential expression of CD14 and CD16 (**B**). Bar chart showing cell fractions of all studied innate immune cell populations before (pre) and 60 minutes after the short intervention (**C**).

**Table 1.** Demographic data.

|  | n | Age mean, SD | Female n (%) | BMI, SD | Daily<br>cigarettes | Years<br>smoking |
| --- | --- | --- | --- | --- | --- | --- |
| Controls | 50 | 47.24 ± 17 | 35 (70) | 23.4 ± 2.7 | NA | NA |
| Smokers | 26 | 41.77 ± 12 | 19 (73) | 24.5 ± 2.5 | 7.9 ± 5.6 | 16 ± 5.6 |
| Comparison (p-value) |  | 0.1696 | 0.156 | 0.1053 |  |  |

### Whole blood lysis and staining

Blood was sampled from the antecubital area via peripheral venipuncture into a sterile BD Ethylenediaminetetraacetic acid (EDTA) Vacutainer blood collection tube with 1.8mg EDTA per milliliter of blood. All samples were processed within 30 minutes to 1 hour. One hundred μL of whole blood was lysed with 1X Red Blood Cell Lysis Buffer (Bio Legend, 10X) following the manufacturer’s instructions. After lysis, cells were centrifuged at 400*g* for 5 min at 18°C. Supernatant was discarded and cells in the pellet were resuspended in 4mL PBS with the aforementioned settings. Cells were further resuspended in 300μL FACS Buffer (1X PBS, 2-5 % FBS, 2 mM EDTA and 2 mM NaN3) and 100µl were added to a 96-well plate to continue with the staining protocol. To avoid nonspecific binding of antibodies to Fc-receptors, samples were incubated for 10 min in 5µl of Human TruStain FcX (BioLegend). Each sample was diluted, divided for 2 panels and stained with the following conjugated antibodies: anti-human CD16 (FITC), anti-human HLA-DR (Peridinin chlorophyll protein-Cyanine5.5), anti-human CD86 (Allophycocyanin), anti-human CD3 (Alexa Fluor 700), anti-human CD66b (Alexa Fluor 700), anti-human CD19 (Alexa Fluor 700), anti-human CD56 (Alexa Fluor 700), anti-human CD36 (Allophycocyanin-Cyanine 7), anti-human CD163 (Brilliant Violet 421), anti-human CD15 (Brilliant Violet 510), anti-human HLA-ABC (Brilliant Violet 605), anti-human CCR2 (Phycoerythrin), anti-human CD62L (Phycoerythrin -Dazzle 594), anti-human CD15 (Phycoerythrin -Cyanine5), anti-human CX3CR1 (Phycoerythrin-Cy7), anti-human CD14 (Alexa Fluor 700), anti-human CD15 (APC), anti-human CD4 (Allophycocyanin-Cyanine 7), anti-human HLA-DR (Brilliant Violet 421), anti-human CD3 (Brilliant Violet 510), anti-human CD19 (Brilliant Violet 605), anti-human CD16 (Brilliant Violet711), anti-human CD45 (FITC), anti-human CD56 (Phycoerythrin), CD8 (Phycoerythrin-Dazzle 594) and anti-human CD20 (Peridinin chlorophyll protein-Cyanine5.5). After an incubation period of 30 minutes at 4°C, samples were washed twice, and were resuspended in 210µL of FACS-Buffer for further processing. Fluorescence minus one (FMO) controls were used to determine and assess background fluorescence in the respective detection channel. Data were acquired on the Attune NxT Flow Cytometer (Thermo Fisher Scientific) and analyzed with Flowjo Analysis Software (v10.5.3). The focus of our analysis consisted of the single cell selection and characterization of monocytes (HLA-DR^+^ CD3^-^ CD19^-^ CD56^-^ CD66b^-^). Thereafter, cells were further divided in subsets and identified as classical (CD14^++^CD16^-^), intermediate (CD14^++^CD16^+^), and nonclassical (CD14^+^CD16^++^) monocytes (Figure 1B) (Guilliams et al. 2018). Analysis of the frequency of all subsets is shown in Figure 1C. Moreover, neutrophils were characterized as CD15^+^ CD16^+^ and additionally subdivided based on their expression of CD62L into mature (CD16^bright^CD62L^bright^), banded (CD16^dim^CD62L^bright^) and hyper segmented (CD16^bright^CD62L^dim^) neutrophils (van Staveren et al. 2018) (Figure 3B).

**Figure 2.**
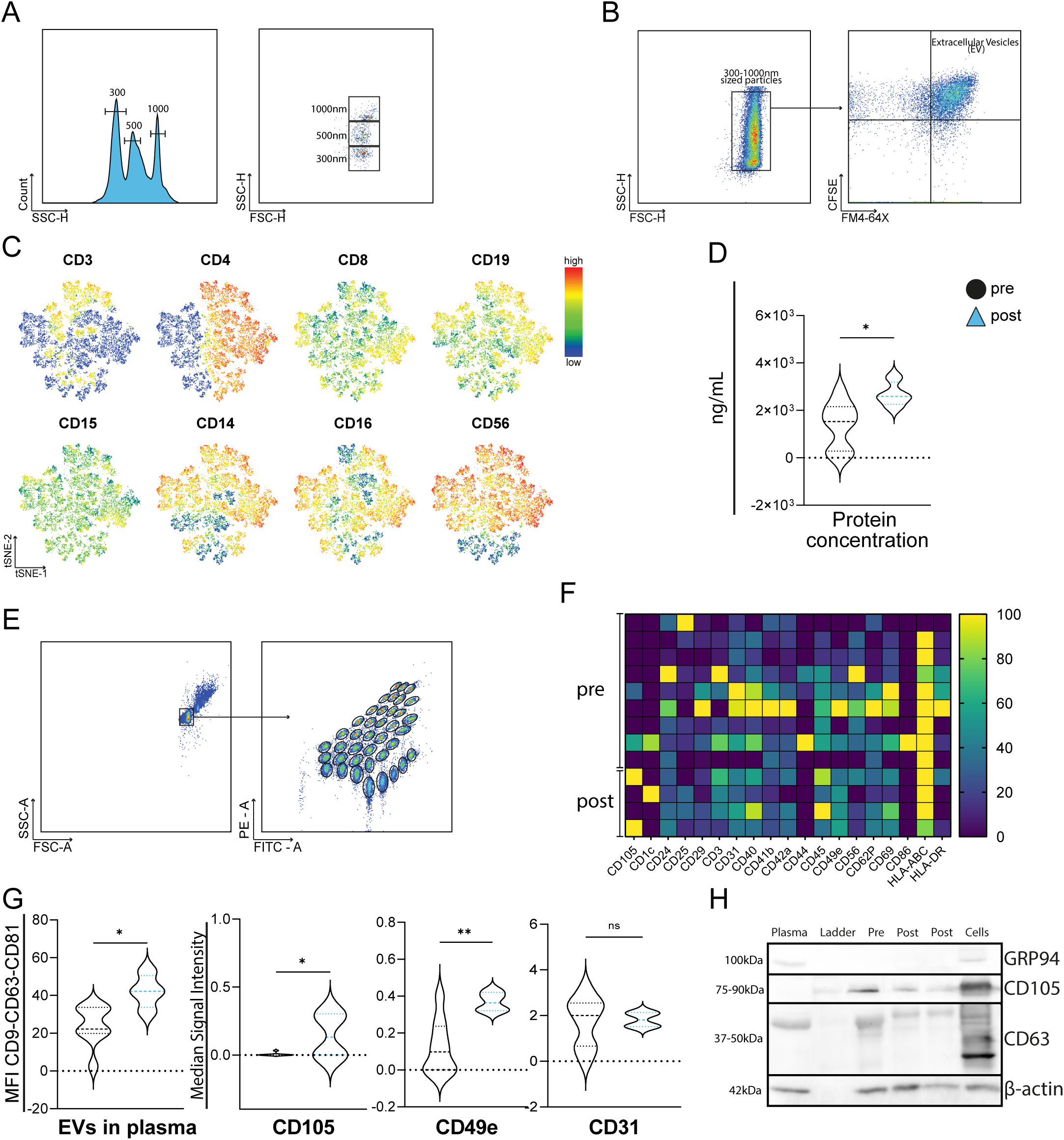
Plasma extracellular vesicles increase after one tobacco smoking session expressed by immune and endothelial cells. Identification of red fluorescent silica beads via flow cytometry. A size range from 300 to 1,000 nm was applied for size calibration (**A**). EV pellets were labeled by protein-specific fluorescent dye (CFSE), and by lipid-specific (FM 4-64FX) dye prior to measurements. Events within the double positive gate were identified as microvesicles (**B**). t-distributed stochastic neighbor embedding (t-SNE) plots of the mean fluorescence intensity of CD3, CD4, CD8, CD19, CD15, CD14, CD16 and CD56 surface antigens in a total of 50,000 double positive events (**C**). Floating bar charts showing the concentration of protein before and after the tobacco smoking intervention, measured via Bradford assay (**D**). Representative gating strategy of the MACSPlex exosome assay, showing the bead size selection (right) and the 37 different dye-labeled capture beads against 37 different EV surface antigens based on FITC and PE fluorescence (left) (**E**). Heatmap of CD9-CD63-CD81 normalized MFI in all individual samples, showing 19 differentially expressed surface markers in our cohort both before and after the intervention (**F**). Violin plots showing the mean MFI of the three main EV markers CD9, CD63 and CD81 in each cohort (far left) and the signal intensity of CD105, CD49e and CD31 surface antigens after detection of CD9, CD63 and CD81 (**G**). Presence of characteristic EV (CD63), plasma (β-actin), endothelial (CD105) and cell lysate (GRP94) markers verified by western blotting before (pre), and 60 minutes after smoking (post) (**H**). All statistical evaluations were performed using non-parametric unpaired t-test. P values: *≤ 0.05; ** for p≤ 0.001; *** for p ≤ 0.0001.

**Figure 3.**
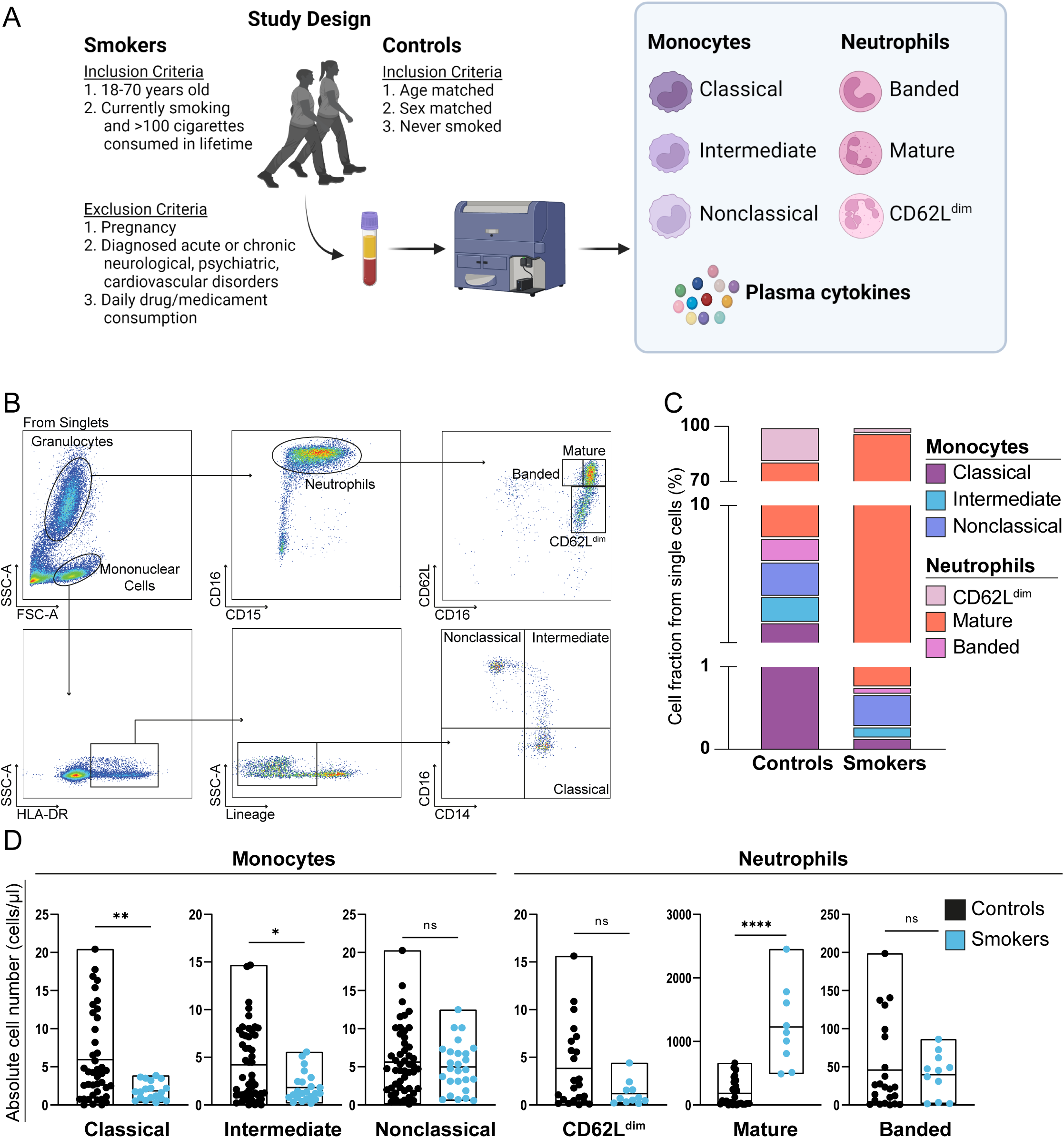
Circulating monocyte and neutrophil subsets are altered in smokers compared to nonsmoking healthy controls. Clinical and experimental study design and analytical processes showing inclusion and exclusion criteria for healthy controls and smokers groups. Peripheral blood was withdrawn and further studied via Flow Cytometry to analyze innate immune system cell subsets and circulating cytokines (**A**). Representative gating strategy of flow cytometry data. Single cells were selected first (not shown), afterwards granulocytes and mononuclear cells were gated based on their size and granularity. Neutrophils were gated based on their expression of CD16 and CD15, subpopulations were gated using CD62L. Mononuclear cells positive for HLA-DR (MHC-II), and negative for lineage markers (CD3, CD19, CD56 and CD66b) were then studied and further subdivision in classical, intermediate and nonclassical monocytes based on their expression of CD14 and CD16 (**B**). Mekko chart representing the cell fraction contribution of all studied innate immune cell populations in controls versus smokers (**C**). Floating bar charts showing the absolute cell numbers of monocytes and neutrophils subsets in controls and smokers (**D**). Statistical evaluations were performed using non-parametric unpaired t-test. P values: *≤ 0.05; ** for p≤0.001; *** for p ≤ 0.0001.

### Extracellular vesicles isolation

Blood from the participants (n=13) were collected on ACD tubes to prevent in vitro vesiculation of platelets and blood cells (György et al. 2014). Medium sized vesicles (microvesicles) were separated by differential centrifugation as described previously (Pospichalova et al. 2015). Briefly, 2 mL of platelet-free plasma were first centrifuged at 200*g* for 5 min to remove live cells. Thereafter, supernatants were transferred into new sterile tubes and further centrifuged at 1,500*g* for 10 min for the clearance of large EVs. To extract microvesicles, supernatants were transferred once more for ultracentrifugation at 14,000*g* for 70 minutes at 4°C. The supernatant was discarded, and the pellet containing microvesicles was resuspended in 0.22µm (pore size filter) filtered PBS. This last ultracentrifugation step was repeated to improve microvesicle enrichment and purity. Remaining pellets containing microvesicles were stored in 200µl of filtered PBS at −80°C until further processing. The guidelines of the International Society for Extracellular Vesicles (Théry et al. 2018) recommend referring to EVs by their physical or molecular characteristics given that they vary in size, and as yet there are no established markers to identify different by their biogenetic routes. Here, we use the term “mEVs” where we refer to medium sizes nanoparticles

### Extracellular vesicle protein concentration

To determine total protein content, aliquots of frozen mEVs were thawed at 37°C and Bio-Rad Protein Assay (Bradford 1976) was performed following manufacturer’s instructions (500-0006 Bio-Rad. München, Germany). Samples were diluted with filtered PBS 1:50 and bovine serum albumin (BioRad Laboratories, USA) was used as standard (1 mg/mL). Two hundred µL of the diluted sample and 50μL dye (1x Bradford reagent) were added to a 96-well plate and the absorbance was measured at 595 nm using the SpectraMax® M5 Microplate Reader (Molecular Devices LLC).

### Extracellular Vesicle Flow Cytometry

For the analysis of fluorescently labelled microvesicles via flow cytometry, aliquots of frozen mEVs were thawed up at 37°C and centrifuged for 10 min at 14,000*g* to remove aggregates. Supernatant was removed with caution and pellets were resuspended 200μL filtered PBS. Hundred μL per sample were collected separately for technical replicates and staining controls. mEVs were fluorescently labelled with protein-(Carboxyfluorescein succinimidyl ester, CFSE) and lipid-(FM) specific dyes, without the necessity of removing the unbound fluorescent dyes by ultracentrifugation, which further facilitates the characterization of microvesicles for flow cytometry (Pospichalova et al. 2015). Samples were first incubated for 20 min in a final concentration of 100μmol/L CFSE (eBioscience) at 37 °C in the dark. Samples were thereafter incubated for another 10 min at 37 °C in the dark in a final concentration of 20μg/mL FM4-64FX. Antibody labeling was performed subsequent to CFSE/FM4-64FX incubation, samples were stained with anti-human CD3 (Peridinin chlorophyll protein-Cyanine5.5), anti-human CD15 (Allophycocyanin), anti-human CD4 (Alexa Fluor 700), anti-human CD14 (Brilliant Violet 510), anti-human CD19 (Brilliant Violet 605), anti-human CD16 (Brilliant Violet 711), anti-human CD56 (Phycoerythrin) and anti-human CD8 (Phycoerythrin-Dazzle 594). Following incubation, samples were washed, resuspended in filtered PBS and transferred to sterile tubes for data acquisition.

Sample data were acquired on the AttuneNxT Flow Cytometer equipped with 405-, 488-, 561-, and 633-nm lasers and an auto sampler allowing the procession of 96-well plates. To create a stable, slow velocity stream for optimal acquisition recommended for the detection of mEVs, sample collection speed was adjusted to the lowest flow rate (25µL/min) with SSC-threshold set to 0.2 x 10^3^. To exclude carryover of fluorescent events between samples, 10% bleach and filtered PBS washes were included after every sample. Hundred μL of sample suspension was loaded, and a stopping option was set to 50μL. Red fluorescent 300 nm, 500 nm and 1000 nm diameter silica particles (Creative Diagnostics) were used for the correct size identification of medium sized EVs. Size reference was determined through Forward and Side Scatter light (FSC-H and SSC-H). Thereafter, CFSE- and FM 4-64FX-positive events with the designated size were identified as microvesicles. Fluorescence minus one (FMO) and unstained controls were used to aid in the gating strategy (Supplementary Figure 1). Results were exported and analyzed in FlowJo version v10.5.3 (TreeStar). T-distributed stochastic neighbor embedding (tSNE) plots were generated in FlowJo using the DownSample plug-in to normalize the interrogation of events per sample per group. Twenty-five thousand events for each study group were exported and concatenated followed by tSNE algorithm on all compensated parameters for 1,000 iterations, perplexity of 30, learning rate of 4,050, approximate random projection forest – ANNOY as KNN and FFT interpolation as gradient algorithm.

### MACSPlex Exosome kit

To study in more detail the surface markers expressed on our EV population, we performed an in depth study of 37 extracellular vesicles surface epitopes (130-108-813, MACSPlex Human Exosome Kit; Miltenyi, Bergisch Gladbach, Germany) as previously described (Koliha et al. 2016). Briefly, this assay uses phycoerythrin and fluorescein isothiocyanat-labeled polystyrene capture beads that permit the discrimination of 39 bead subpopulations based on their different fluorescence intensities. These beads are added to the plasma-separated mEVs and incubated overnight. The next day, allophycocyanin labeled detection antibodies are added, containing anti-CD9, anti-CD63 and anti-CD81 antibodies that allow for the identification of extracellular vesicles. This process creates a structure consisting of the capture beads, EV and the detection antibodies allowing for the detection of the event and the identification of its surface epitopes. This assay was measured in the Attune NxT flow cytometer collecting the median fluorescence intensity (MFI) of each EV surface marker that was then normalized to the mean MFI of the specific EV markers.

### Western Blot

Western blot analysis was performed on 30µg of isolated mEVs per sample, 0.5µl human plasma and 20µg of human urinary bladder cell lysate (300452CL, CLS, Germany) were used as controls. Samples were thawed at 4°C, total proteins were separated on a gradient sodium dodecyl sulphate-polyacrylamide gel electrophoresis 4% - 10% gel and transferred onto a nitrocellulose membrane. Blocking was performed in 5% non-fat milk and subsequently, incubation overnight with specific primary antibodies at 4°C was performed. The antibodies used were GRP94 (1:1000, 100 kDa, Proteintech, 14700-1-AP, IL, USA), CD105 (1:500, 75-90 kDa, Proteintech, 10862-1-AP, IL, USA), CD63 (1:500, 37-50 kDa, Proteintech, 25682-1-AP, IL, USA) and β-actin (1:1000, 42kDa, Cell Signalling Technology 4970, Cambridge, UK). The following day, membranes were washed with a blocking buffer to remove unbound antibodies, and a host-specific secondary antibody with matching horseradish peroxidase-conjugated substrate was added and incubated before analysis of its chemiluminescence using the ChemiDoc XRS+ System (Bio-Rad, CA, USA).

### Bead-based Cytokine Assay

Plasma samples were thawed at 4°C and centrifuged at 400*g* for 10 min at 4°C to remove debris. Plasma levels of cytokines IL-18, IL-1β, IL-10, TNF-α, IL-8, IL-33, and of soluble triggering receptor expressed on myeloid cells 2 (sTREM2) and brain-derived neurotrophic factor (BDNF) were assessed using the Human LEGENDplex Multiplex Assay (BioLegend®) was used following manufacturer’s instructions (Lehmann et al. 2017). Briefly, the assay contains allophycocyanin-coated beads conjugated with surface antibodies allowing for specific binding to the cytokine of our interest. After incubation of the capture beads with the plasma sample, these compounds were now stained with biotinylated detection antibodies that bind specifically to its analyte on the capture beads. Therefore, a capture bead-analyte-detection antibody sandwich is created, and it’s further stained with streptavidin-phycoerythrin. This last staining provides different intensity signals depending on the amount of bound analyte. Samples were measured using the Attune NxT Flow Cytometer. The concentration of each particular analyte is determined on a known standard curve using the LEGENDplex™ Data Analysis Online Software Suit. Half of the limit of detection (LOD) was used to perform statistical analyses when a sample value fell below the LOD.

### Statistical Analysis

All statistical analyses were performed using GraphPad Prism 9. We evaluated two groups in the chronic setting (Smokers and Controls) and two groups on the acute intervention (pre and post intervention). All data were non-normally distributed as shown by D’Agostino & Pearson and Shapiro-Wilk tests, therefore, non-parametric unpaired t-test was performed. Graphical representations of data were performed in Graphpad Prism 9 and BioRender. Our analyses were adjusted for potential confounds that could be associated with blood cell values, such as age, sex and BMI (Table 1). Representative gating strategy of Flow Cytometric data was analyzed and performed using FlowJo software. An alpha level of p≤0.05 was used for all statistical tests. Therefore, p values ≤ 0.05 were considered statistically significant and marked with asterisks as follows: * for p ≤ 0.05; ** for p≤0.001; *** for p ≤ 0.0001.

## Results

### Immune cell populations shift after an acute cigarette smoking intervention in never-smokers

In order to investigate the acute effect of cigarette smoking in healthy nonsmokers, we asked never-smokers to smoke two cigarettes (12 tar yield mg/cig, 1.0 nicotine yield), and we evaluated their innate and adaptive immune cell subset composition before and one hour after the intervention (Figure 1A) via multicolor flow cytometry using the gating strategy shown in Figure 1B. We found no significant changes in the absolute cell number neither of said subsets nor in the cell fractions (Supplementary Table 1). However, a slight increase in the neutrophil (CD15^+^ CD16^+^) and classical monocytes (CD14^++^CD16^-^) fraction was observed after the intervention. Interestingly, a moderate decrease in the CD3^+^ CD4^+^ and CD3^+^ CD8^+^ T, B (CD3^-^ CD56^-^ CD14^-^ CD20^+^), NK (CD3^-^ CD56^+^) and NKT (CD3^+^ CD56^+^) cell fractions was found after the smoking intervention (Figure 1C). These results point towards a plausible acute immunosuppression after the one cigarette smoking session.

### Extracellular vesicle concentration increases in circulation 1 hour after smoking in never-smokers

After the assessment of the immune cell populations, we set to investigate the role of acute smoking in the release of extracellular vesicles. Plasma-derived mEVs were measured before and one hour after smoking. An overall increased concentration of plasma-derived mEVs was found after the acute smoking intervention (Figure 5D) (p=0.0305). This result was confirmed by immunostaining of three canonical tetraspanins found in association with EVs: CD9, CD63 and CD81 (Figure 2G) (p=0.0121). Our findings are supported by previous research where an increased release of EVs was reported from platelets, leukocytes and endothelial cells after tobacco smoking (Mobarrez et al. 2014).

**Figure 4.**
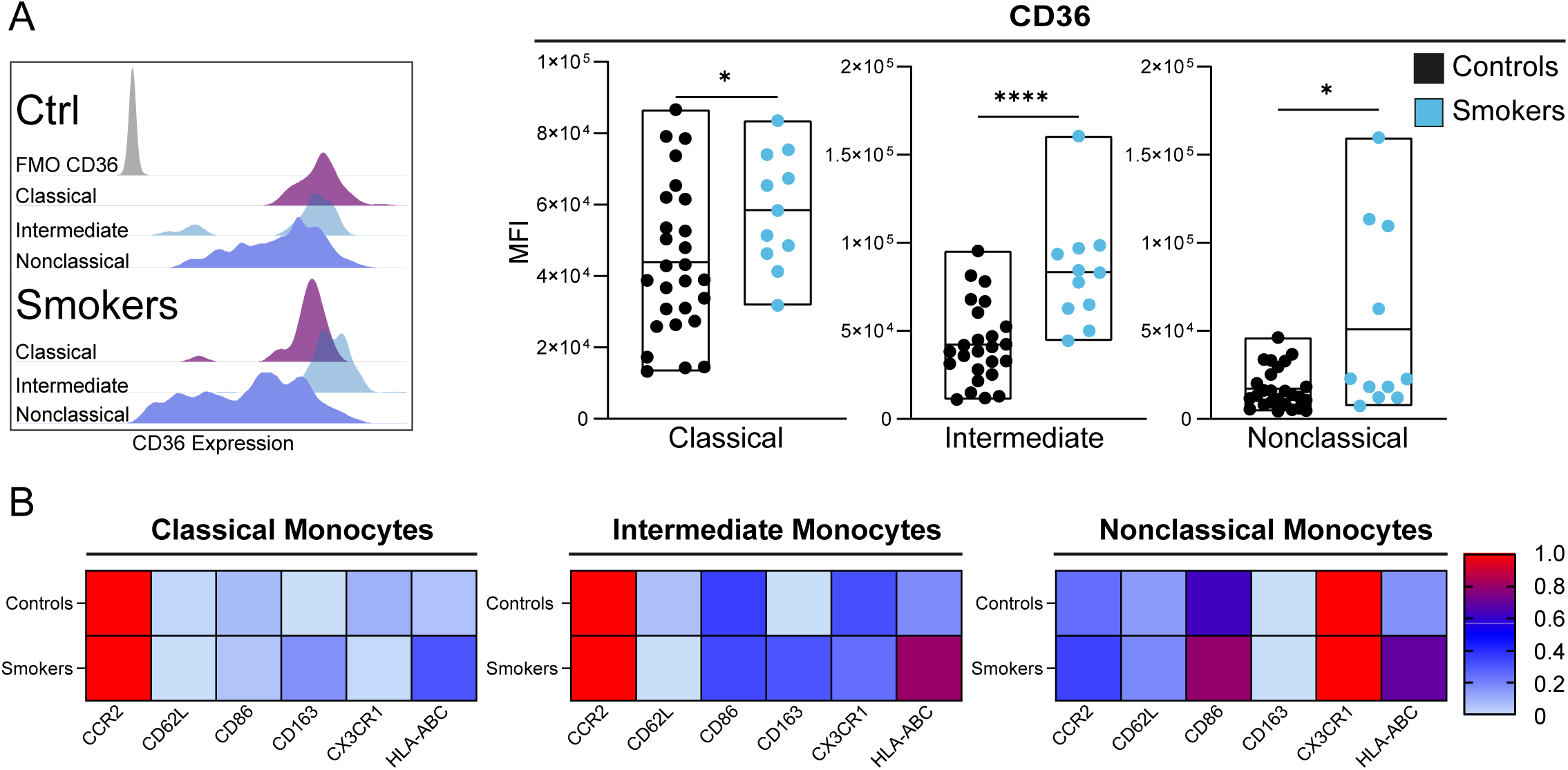
Chronic tobacco smoking alters CD36 expression in the monocyte subsets. Representative histograms of CD36 median fluorescence intensity (MFI) in monocyte subsets in the two cohorts. Floating bar charts depicting MFI of CD36 in all monocyte subsets from smokers compared to controls. (**A**). Heatmaps showing the normalized MFI of key activation markers (CCR2, CD62L, CD86, CD163, CX3CR1 and HLA-ABC) across all monocyte subsets (**B**). Statistical evaluations using non-parametric unpaired t-test. P values: *≤ 0.05; ** for p≤0.001; *** for p ≤ 0.0001

**Figure 5.**
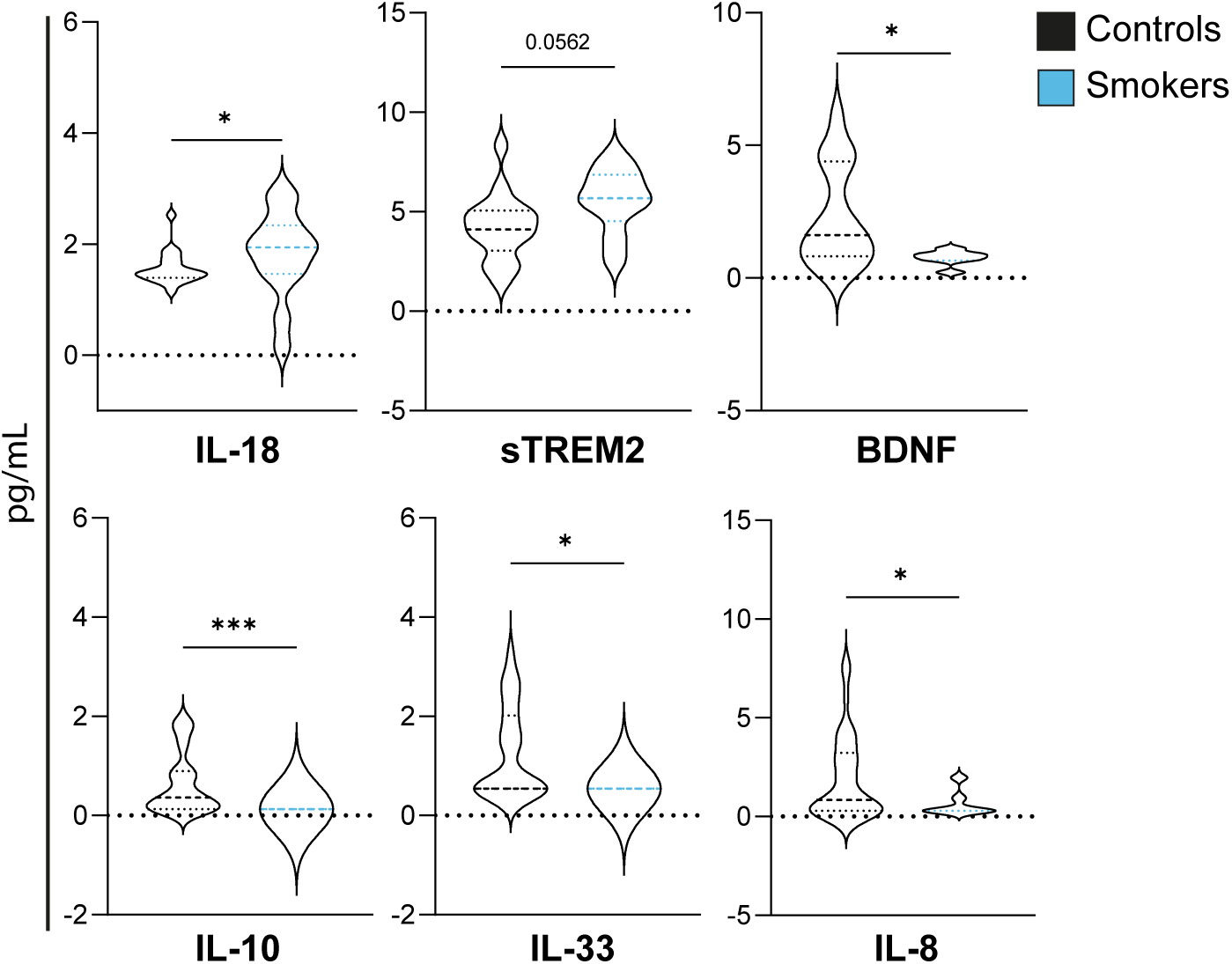
Circulating cytokine changes in chronic smokers. Levels of IL-18, sTREM2, BDNF, IL-10, IL-33 and IL-8 present in plasma detected by flow cytometry. Values are shown in pg/mL, data points below the Limit of Detection (LOD) were calculated as half the LOD. Statistical evaluation using non-parametric unpaired t-test. P values: *≤ 0.05; ** for p≤0.001; *** for p ≤ 0.0001. sTREM2 – soluble Triggering receptor expressed on myeloid cells 2. BDNF – b rain-derived neurotrophic factor.

To assess whether these mEVs were from immune cell origin, we performed a comprehensive study to characterize the EV origin based on their expressed surface markers. We utilized multicolor flow cytometry to classify identify microvesicles and by utilizing fluorescent silica beads, its size was determined. Our gating strategy was defined for nanoparticles sized between 300 nm up to 1000 nm (Figure 2A). Flow cytometry showed predominantly double positive mEVs (Figure 2B) with signal distribution which indicated ≥ 95% mEVs based on the membrane markers CFSE and FM 4-64X (Pospichalova et al. 2015). We further investigated the source of double positive events by using surface markers of the main immune cell populations, namely CD3, CD8 and CD4 for T cells, Cytotoxic T cells and T helper cells respectively; CD19 for B cells, CD15 for neutrophils, CD14 and CD16 for monocytes and CD56 for natural killer cells. Our results showed that the main immune origin of the mEVs after a short cigarette smoking session were monocytes, NK, T, and B cells (Figure 2C). Afterwards, to assess the non-immune cell-derived mEVs, we studied 37 exosomal surface epitopes using a bead-based assay (Figure 2E). Nineteen differently expressed markers between pre and post tobacco smoking intervention were selected (Figure 2F). Out of all the investigated EV surface epitopes in the plasma-isolated mEVs, the endothelial transmembrane glycoprotein Endoglin (CD105) in addition to the endothelial activation marker CD49e (integrin α-5) were significantly increased upon acute tobacco intervention (Figure 2G) (Słomka et al. 2018). In addition, following the MISEV2018 criteria, Western Blot confirmed the EV identity and CD105 presence in plasma-derived mEVs from chronic smokers (Théry et al. 2018). Presence of tetraspanin CD63 was evidenced, along with the absence of Golgi membrane stacking protein GRP94, which was only found in cell lysates (Figure 2H). Taken together, these results point towards an activation of endothelial cells followed by the release of mEVs even after a short tobacco smoking session in healthy nonsmoker individuals.

### Clinically asymptomatic current smokers show a decrease in classical and intermediate monocytes accompanied by an increased number of mature neutrophils

To further understand the role of chronic smoking, we now assessed otherwise healthy chronic smokers (Figure 3). Peripheral monocyte dynamic subsets play an important role in modulating innate immune responses and initiating adaptive immunity (Patel et al. 2017). Here, to expand the understanding on the profile of the peripheral innate immune system of healthy smokers, we studied whole blood from healthy smokers and nonsmokers. No significant differences were observed in the overall monocyte population (p= 0.1921) however, we found the classical and intermediate monocyte subsets to be significantly decreased in chronic smokers (p=0.0042 and p=0.0284 respectively). No differences were found within the nonclassical monocytes after comparing never smokers versus chronic smokers (p=0.8469) (Figure 3D). We found non-significant lower levels of the CD62L^dim^ and CD62L^bright^ subsets (p=0.5134 and p=0.1162, respectively) and interestingly, a prominent higher fraction of the mature neutrophils was evidenced in the chronic smokers when compared to nonsmoker controls (p=<0.0001) (Figure 3D). These results indicate a significantly different innate immune population composition in current smokers is already present.

### Increased CD36 expression by monocytes in asymptomatic chronic smokers

Given the contribution of CD36 to the development of dysfunctional endothelium and its role in atherosclerotic lesions, we focused our analysis of its expression on all monocyte subsets. CD36 is a class B scavenger receptor present on an extensive range of cells including monocytes, macrophages, microvascular endothelial cells and adipocytes (Park 2014). However, due to our interest on the peripheral innate immunity and the practicality of a liquid biopsy, we focused our study on the early role of monocyte subsets by analyzing the expression of surface marker CD36 per subset (Figure 4A). Interestingly, higher levels of CD36 were a general feature detected on all monocyte subsets in chronic smokers (p=0.0453, p=>0.0001), and p=0.0469, respectively). CD36 mediates foam cell formation and promotes atherosclerosis (Tian et al. 2020). In line with previous literature, we found an overall higher expression of CCR2 in classical (CD14^++^CD16^-^) and intermediate (CD14^++^CD16^+^) monocytes, which was then decreased in nonclassical monocytes (CD14^+^CD16^++^). Similarly, CX3CR1 was found highly expressed in nonclassical monocytes (CD14^+^CD16^++^) and lessened in (CD14^++^CD16^-^) and intermediate (CD14^++^CD16^+^) monocytes (Kapellos et al. 2019; Hristov and Heine 2015). No significant differences were found in levels of CD62L, CD86, CD163, or HLA-ABC when comparing asymptomatic chronic smokers with never-smokers. (Figure 4B). These results show that the expression of CD36 on circulating monocytes is increased in chronic smokers, suggesting an early shift towards a pro-atherosclerotic environment in otherwise healthy individuals with history of smoking in the absence of clinical symptoms.

### Chronic tobacco smoking in healthy individuals influences peripheral cytokine levels

After the observation of highly activated monocyte subsets, we aimed to test whether plasma cytokine levels would be also altered. Diverse cytokines have been already studied in chronic smokers, particularly pro-inflammatory cytokines such as IL-6 and IL-1β levels, which have been found increased (Elisia et al. 2020; Barbieri et al. 2011). Therefore, our focus was directed to the investigation of IL-18 levels, due to its close link with IL-1β as byproducts of the NLPR3 inflammasome activation. NLPR3 inflammasome has been found to be highly active in aortic endothelial cells after nicotine treatment *in vitro* consecutively leading to an increased production of IL-18 (Wu et al. 2018). Similarly, IL-18 has been associated with higher risk for coronary heart disease and tobacco smoking (Jefferis et al. 2011). Our results revealed a significant increase of circulating IL-18 in asymptomatic chronic smokers (p=0.0396) that was accompanied by a decrease in anti-inflammatory cytokine IL-10 (p=0.0007) (Figure 5). Among with these changes, another member of the IL-1 family of cytokines, the alarmin IL33, was found decreased (p=0.0203) in the chronic smokers groups compared to never smokers (Liew et al. 2016). IL-8 is a pro-inflammatory cytokine that has been described to participate in atherosclerotic plaque destabilization and as a mortality predictor in patients with acute coronary syndrome (Cavusoglu et al. 2015; Boekholdt et al. 2004). In the current smokers’ cohort however, the plasma levels of this cytokine were also found to be decreased (p=0.0430).

All previous results from chronic smokers have pointed towards peripheral ‘silenced’ inflammation. It has been described that peripheral inflammation, evidenced by activation of the innate immune system and release of proinflammatory cytokines can affect the brain (Huang et al. 2021). For this reason, we assessed BDNF and sTREM2 levels in plasma to investigate whether the long exposure to tobacco smoking altered soluble levels of these markers. Our results show significantly lower levels of BDNF in current smokers, similarly to previous literature (Bhang et al. 2010), accompanied by a trend of higher soluble TREM2 in plasma of asymptomatic chronic smokers (p=0.0562).

## Discussion

Long term tobacco smoking has been strongly linked to the development of various cardiovascular and neurological disorders, with vascular endothelial damage constituting an important pathomechanism in the development of hypertension and atherosclerosis (Messner and Bernhard 2014). After the onset of cardiovascular disorders, chronic smokers endure a higher risk for developing vascular dementia due to the damage on the endothelium (O’Brien and Thomas 2015). These changes take place after the persistent and chronic stressor that smoking represents. However, the acute effects of smoking on the vasculature and the long-term effects in asymptomatic chronic smokers remained largely unmapped.

In the present study, we first investigated the early and ongoing effects of smoking on the immune and vascular fitness in healthy volunteers. One focus was on the initial effects of smoking on the blood immune cell compartment and early endothelial damage in a healthy organism that has not been exposed consistently to tobacco smoking before. Our results reveal that already a very short acute tobacco smoking challenge influences the composition of innate and adaptive immune cells in the blood of the volunteers. Activated cells can signal through EVs, small membrane-enclosed vesicles that are released by every cell in the organism and can be detected in body fluids (Alberro et al. 2021; Panteleev et al. 2017; Buzas et al. 2014). Immune and non-immune cells did release an increased amount of EVs, after the acute tobacco smoking intervention. These data are in line with previous literature in which the amount of EVs (microparticles, an old designation for mEVs) was increased after cigarette smoking (Mobarrez et al. 2014; Heiss et al. 2008).

Next, we investigated the source of mEVs after a single smoking session, and our results revealed that the increased levels of mEVs were largely of endothelial origin. The elevated number of mEVs after acute smoking was previously observed in airway basal cells responding to CSE *in vitro* (Saxena et al. 2021). The vascular endothelium is comprised of a monolayer of endothelial cells as the first barrier of the vessel wall, structurally supporting the circulatory system, and as a versatile endocrine organ producing of different molecules (Baumgartner-Parzer and Waldhäusl 2001; Osteikoetxea et al. 2016). In response to smoke exposure, endothelial cells can release inflammatory and proatherogenic cytokines and reactive oxygen species (ROS), leading to endothelial dysfunction and damage. This has been described in chronic smokers, although the acute *in vivo* effects of smoking in healthy individuals have not been fully elucidated. In 2004, Papamichael et al reported that healthy individuals who smoked one cigarette had a decrease in flow mediated dilatation (FMD), suggesting a transient endothelium dysfunction from 15 minutes to an hour after the intervention (Papamichael et al. 2004). Our results indicate that the endothelium secreted a sizable number of mEVs, characterized by the expression of CD105 and CD49e on the surface of isolated mEVs. CD105 is a transmembrane protein and co-receptor for transforming growth factor β (TGF-β) expressed on the surface of endothelial cells, it has been recognized as a marker of angiogenesis, extravasation of leukocytes and associated with tissue injury (Cheifetz et al. 1992; Duff et al. 2003; Rossi et al. 2019). On the other hand, CD49e is the integrin subunit alpha 5 expressed on endothelial cells. The absence of integrin α5 results in leakage of the blood-retinal barrier in mice, highlighting its critical role in barrier integrity (Ayloo et al. 2022).

It has been long known that tobacco smoking has strong negative impact on vascular, cardiac, pulmonary and neurological health, including high toxicity that can lead to different types of cancer, chronic obstructive pulmonary disease and vascular dementia, among other disorders (Jha et al. 2013; Le Foll et al. 2022; U.S. Department of Health and Human Services). Previous studies have shown elevated levels of circulating leukocytes in blood of chronic smokers. However; the focus of these studies is predominantly on patients with already arterial hypertension, COPD or lung cancer (Andersson et al. 2019; Elisia et al. 2020; Gordon et al. 2011).

The first innate immune response after a stressor is guided by neutrophils (Hidalgo et al. 2019). Our results reveal elevated neutrophil numbers, which can be mechanistically explained by the immediate innate immune response towards the insult that represents tobacco smoking. It has been described however, that even though high levels of neutrophils are found in chronic smokers, their function may be compromised and reprogrammed and, as a result of tobacco smoking, may predispose the host to the development of chronic diseases and impair their response against infections (Zhang et al. 2018; Valiathan et al. 2014; Aghapour et al. 2022). Next, we focused on monocyte subsets given their critical role in the development of atherosclerosis, driving disease progression via their recruitment into atherosclerotic plaques. We observed significantly lower frequencies of both classical (CD14^++^CD16^-^) and intermediate (CD14^++^CD16^+^) circulating monocytes in chronic smokers. Our findings suggest an early monocyte suppression upon chronic tobacco smoking that in later stages yields higher levels of all innate immune system cells promoting clinical symptomatology. It could also describe a transient adaptation of monocytes due to the chronicity of the stressor. Of note, our study’s cohort include healthy nonsmokers, whereas in previous literature, chronic smokers with symptoms and/or diagnoses of tobacco and non-tobacco related diseases such as COPD, cancer, ischemic heart disease, arterial hypertension and diabetes mellitus were present in a relevant fraction of the participants included (Pedersen et al. 2019; Elisia et al. 2020; Gordon et al. 2011).

Next, we characterized the activation of monocyte subsets especially focusing on CD36, a scavenger receptor heavily involved in atherosclerotic plaque formation. CD36 is a scavenger receptor that mediates oxidized LDL (oxLDL) uptake and its expression contributes to macrophage foam cell formation (Febbraio et al. 2001). Besides its role in lipid metabolism, it has also been described to be involved in angiogenesis, atherosclerosis and inflammation (Tian et al. 2020). *In vitro* studies have shown that CD36 depletion from monocytes and macrophages profoundly affect the ability of uptake oxLDL and prevented the formation of foam cells, which are crucial for the development and progression of atherosclerosis (Febbraio et al. 2001; Park 2014). In our study, the expression of CD36 was significantly higher on the surface of classical, intermediate and nonclassical monocytes, showing that even though the monocyte frequency was not affected, chronic smoking induces the excessive upregulation of CD36. This promotes tissue damage even before the onset of any clinical symptomatology, and a myeloid immune specific profile can be observed before tobacco-related disease onset. A study by Mehta and Dhawan found that THP-1 cells exposed to cigarette smoke extract (CSE) *in vitro* not only were more activated evidenced by their increased expression of CD36, but additionally they showed an increased activation of the NLRP3 inflammasome accompanied by an increased production of proinflammatory cytokines such as IL-1β and IL-18 (Mehta and Dhawan 2020). Our results showed an increased activation of monocytes based on their highly increased surface expression of CD36.

Further, we measured an increased presence of IL-18 in plasma, which confirms the previous *in vitro* findings indicating an important effect of cigarette smoking on inflammasome-caspase axis evidenced by elevated levels of IL-18 and caspases (Kang et al. 2007; Eltom et al. 2014). In addition, we detected a decrease in IL-10 a broadly expressed anti-inflammatory cytokine (Saraiva and O’Garra 2010). This finding has also been described in the sputum of patients with COPD (Maneechotesuwan et al. 2013), along with findings both *in vitro* and *in vivo* in mice exposed to cigarette smoker or CSE (Higaki et al. 2015).

IL-33, is an alarmin normally released by damaged barrier cells such as endothelial cells (Liew et al. 2016). One of the particularities of IL-33 is that it acts as a traditional cytokine through a receptor complex but also plays an important role as an intracellular nuclear factor that regulates the transcription of the p65 subunit of the NF-κβ complex, involved in endothelial cell activation (Choi et al. 2012; Ali et al. 2011). This cytokine has been found increased in plasma of patients with COPD (Xia et al. 2015) and decreased in mini-bronchoalveolar lavage (mini-BAL) from chronic smokers. A study found increased intracellular mRNA levels of IL-33 in bronchial epithelial cells after induction with CSE *in vitro* (Pace et al. 2014). IL-33 is known for its proinflammatory effect, however in ApoE -/- mice, IL-33 helped reduce the presence of atherosclerotic plaques (Miller et al. 2008). In our study, decreased levels of IL-33 were detected in the plasma of chronic asymptomatic smokers, suggesting an IL-33 intracellular arrest due to its role as a transcription factor. These low levels of IL-33 can potentially be a prognostic factor given that its absence supports the formation of atherosclerotic plaques in healthy humans with history of smoking. Taking together, these cytokine levels suggest a compromised response from the organism against the chronic inflammatory stimulus that represents tobacco smoking.

Smoking has been listed as one of twelve modifiable factors for dementia and, through its detrimental effect on endothelial structure and function, particularly increases the risk for vascular dementia, the second most common cause of dementia after Alzheimer’s disease (Cipollini et al. 2019). The effects by smoking that cause dementia varies from toxic to vascular damage (Livingston et al. 2020). It has been widely described that chronic endothelial damage leads to blood-brain barrier dysfunction and consequent neuroinflammation (Wardlaw et al. 2017; Rajeev et al. 2022). There is a long list of biomarker candidates being considered; however, knowing the role that chronic cigarette smoking has in the presentation of vascular dementia, we aimed to study the plasma levels of soluble triggering receptor expressed on myeloid cells 2 (sTREM2) and brain-derived neurotrophic factor (BDNF) in asymptomatic chronic smokers. sTREM2 is a receptor expressed by activated microglial cells, and it has been described as a reliable neurodegeneration biomarker both in CSF and blood of AD patients and in pre-symptomatic individuals with AD-related Tau pathology (Brosseron et al. 2022). In our study, we measured elevated sTREM2 in chronic smokers, suggesting neuroinflammatory alterations that can be attributed to microglia dysfunction during the course of chronic smoking.

BDNF is the most prevalent growth factor in the brain regulating neuronal plasticity and survival (Pan et al. 1998; Klein et al. 2011; Zuccato and Cattaneo 2009). It is of relevance to recognize that unlike serum BDNF, plasma BDNF is not affected by platelet-stored BDNF due to the absence of a clotting process, and this measurement allows for prediction of BDNF levels in the CNS without an invasive procedure (Gejl et al. 2019; Fujimura et al. 2002). We detected lower levels of peripheral BDNF in plasma of chronic tobacco smokers when compared to never smokers, similarly to previously reported literature (Bhang et al. 2010). This finding goes in line with a large body of evidence suggesting that reduced BDNF availability upon exposure to chronic stressors increases the risk for accelerated aging and also for the development of neurodegenerative and cardiovascular diseases (Miranda et al. 2019; Kaess et al. 2015). Together with the observation that stress-dependent BDNF suppression may be dependent on neuroinflammatory signaling the reduction of BDNF levels accompanied by a trend of increased sTREM2 levels in our study is suggestive for a pre-clinical impact of tobacco smoking on CNS function in clinically healthy individuals (Schott et al. 2021).

Collectively, we demonstrate the effects of a single tobacco smoking session on the endothelial signaling in healthy never-smokers. Even though the characterization of innate immune cell subsets has been attempted previously in smokers, to our knowledge, this is the first study that provides an in-depth characterization of monocyte subsets and their activation profile comparing disease-free and pre-clinical asymptomatic chronic smokers. Furthermore, we provide evidence for altered circulating cytokine levels as well as proteins related to CNS pathology in clinically asymptomatic chronic smokers. These findings provide further insight into the effects of tobacco smoking on the organism and the potential vascular damage that can contribute to neurodegenerative disorders, specifically cerebrovascular dysfunction which lowers the threshold for dementia. This might be of clinical value, due to a possible classification of smokers into high- vs. low- risk groups, guiding their motivation to quit tobacco consumption, and elucidate potential biomarkers to diagnose vascular disorders before symptom onset, allowing for prevention and clinical intervention to improve prognosis (O’Brien and Thomas 2015; Sachdev et al. 2014). In conclusion, we propose an immuno-vascular dysregulated profile in chronic tobacco smokers that are otherwise healthy individuals, which may contribute to the pathogenesis of early preclinical stages of, particularly vascular, dementia

### Strengths and Limitations

The strengths of our study include a comparison between two different disease-free asymptomatic groups, and the in-depth characterization of monocyte subsets and their activation, accompanied by peripheral cytokine levels and extracellular vesicles with one liquid biopsy. Our study also included older adults, who are often excluded from this type of research. A limitation was that the majority of our participants were of European ancestry and therefore it is impossible to extrapolate our results to other ethnicities. In addition, the smoking status of the participants was self-reported and we did not conduct any cotinine or nicotine blood assessment.

## Acknowledgments

This research was supported by DGM Sc 26/1, RTG2413 Synage. A.P.G., B.H.S., and I.R.D. further received support from the European Fund for Regional Development and the State of Saxony-Anhalt (Research Alliance “Autonomy in Old Age”). The project has also received funding from the EU’s Horizon 2020 research and innovation program under grant agreement No. 739593 and by the Hungarian Thematic Excellence Program No. TKP2020-NKA-26 and the National Cardiovascular Laboratory program.

## Author contribution

IRD conceived the presented idea, designed the study, supervised analyses and edited the manuscript. IRD, SS and BHS were responsible for supervision and project administration. EIB and EP supervised the isolation and processing of plasma-derived mEVs and edited this manuscript. APG planned and carried out the experiments, data analyses, data visualization and wrote the manuscript. LM contributed to sample preparation, measurements, analyses and wrote the manuscript. APG, IRD, SS, BHS and LM contributed to the interpretation of the data. All authors approved the submitted version.

---

**Supplementary Figure 1.**
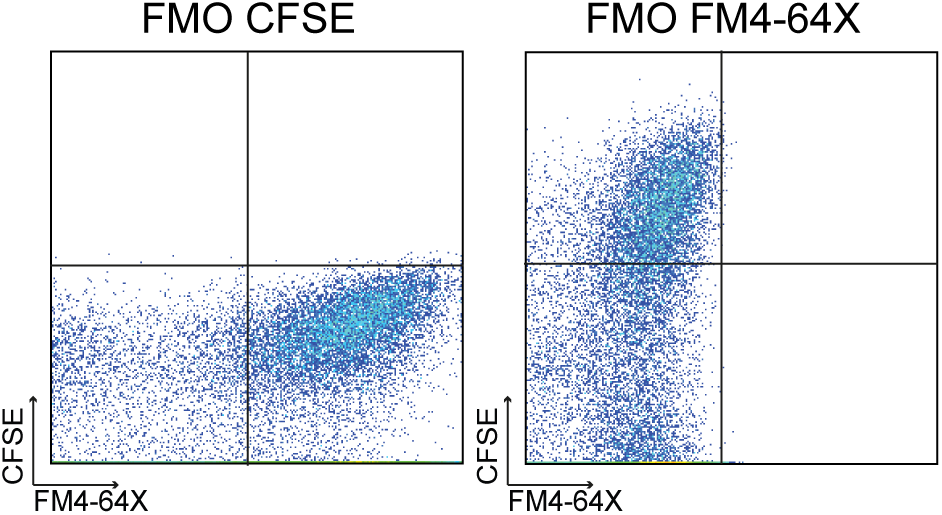
CFSE and FM4-64X Fluorescence Minus One (FMO) Controls. Fluorescence Minus One (FMO) Controls for protein-specific fluorescent dye (CFSE) and lipid-specific (FM 4-64FX) dye prior to measurements. Events within the double positive gate were identified as microvesicles as shown in Figure 2B.

**Supplementary Table 1.** Absolute cell counts.

| | *pre*<br>events/μL<br>(Mean, SEM) | *post*<br>(Mean, SEM) | Comparison<br>(p-value) |
| --- | --- | --- | --- |
| *Classical Monocytes* | 11.11 ± 2.43 | 18.67 ± 5.21 | 0.1261 |
| *Intermediate Monocytes* | 1.481 ± 0.30 | 2.115 ± 0.79 | 0.6094 |
| *Nonclassical Monocytes* | 5.521 ± 0.88 | 7.081 ± 2.04 | 0.6358 |
| *Neutrophils* | 187 ± 38.8 | 279.7 ± 78.04 | 0.4009 |
| *B cells* | 92.96 ± 2.99 | 53.71 ± 4.83 | 0.9009 |
| *NKT cells* | 26.38 ± 7.71 | 35.01 ± 5.23 | 0.0762 |
| *T helpers* | 17.59 ± 15.83 | 17.55 ± 14.37 | 0.3242 |
| *Cytotoxic T cells* | 39.46 ± 4.94 | 31.31 ± 10.85 | 0.5845 |
| *NK cells* | 39.57 ± 6.61 | 17.09 ± 12.16 | 0.2785 |

